# Benchmarking Software for DDA-PASEF Immunopeptidomics

**DOI:** 10.1101/2025.05.28.656277

**Authors:** Yannic Chen, Annica Preikschat, Annette Arnold, Riccardo Pecori, David Gomez-Zepeda, Stefan Tenzer

## Abstract

Mass spectrometry (MS) is the method of choice for high-throughput identification of immunopeptides, which are generated by intracellular proteases, unlike proteomics peptides that are typically derived from trypsin-digested proteins. Therefore, the searching space for immunopeptides is not limited by proteolytic specificity, requiring more sophisticated software algorithms to handle the increased complexity. Despite the widespread use of MS in immunopeptidomics, there is a lack of systematic evaluation of data processing software, making it challenging to identify the optimal solution. In this study, we provide a comprehensive benchmarking of the most widespread/used data-dependent acquisition (DDA)-based software platforms for immunopeptidomics: MaxQuant, FragPipe, PEAKS and MHCquant. The evaluation was conducted using data obtained from the JY cell line using the Thunder-DDA-PASEF method. We assessed each software’s ability to identify immunopeptides and compared their identification confidence. Additionally, we examined potential biases in the results and tested the impact of database size on immunopeptide identification efficiency. Our findings demonstrate that all software platforms successfully identify the most prominent subset of immunopeptides with 1% false discovery rate (FDR) control, achieving medium to high identification confidence correlations. The largest number of immunopeptides were identified using the commercial PEAKS software, which is closely followed by FragPipe, making it a viable non-commercial alternative. However, we observed that larger database sizes negatively impacted the performance of some software platforms more than others. These results provide valuable insights into the strengths and limitations of current MS data processing tools for immunopeptidomics, supporting the immunopeptidomics/MS community in determining the right choice of software.

## Introduction

The Major Histocompatibility Complex (MHC) plays a crucial role in the adaptive immune response by presenting immunopeptides to T-cells, enabling the identification of aberrant cancerous and pathogenic non-self-peptides. MHC are classified into class 1 and class 2, with the former presenting primarily endogenous peptides derived from intracellular proteins while the latter primarily presents exogenous peptides acquired through endocytosis [1][2]. In humans, this protein is termed human leukocyte antigen (HLA) and typically presents peptides of length 8-13 amino acids for class 1 and 13-25 for class 2 [3][4]. The study of the whole set of immunopeptides, known as the immunopeptidome, is fundamental to many medical research fields, particularly immunotherapy, where disease-specific immunopeptides can be leveraged for targeted treatment [1][5][6].

Mass spectrometry (MS) is the method of choice for high-throughput identification of immunopeptides[7]. MS techniques infer peptide sequences by comparing observed spectra to hypothetical peptide fragmentation pattern. However, this approach faces challenges due to missing fragmentation from incomplete fragmentations as well as interfering peaks from contaminants/other biological sources in case of multiplexes spectra. To address this, database search algorithms are employed to make informed predictions of peptide identities by limiting the possible identities to known peptide sequences [8]. It is important to note that peptides may undergo post-translational modifications (PTMs), which alter the fragment masses and expand the range of potential identities. To account for false positives inherent in this approach, False Discovery Rate (FDR) analysis is used as a quality control measure. FDR employs “decoy” databases containing sequences not present in the sample of interest, to assess the reliability of identifications by quantifying the rate of false “decoy hits” [9].

Immunopeptidomics poses unique challenges for database search engines compared to traditional trypsin-based bottom-up proteomics. In contrast to tryptic digests, immunopeptides are generated by intracellular proteases, necessitating searches without restrictions imposed by proteolytic specificities, dramatically expanding the search space and increasing computational demands [1].

Among various modes of measurement, Data-Dependent acquisition mode (DDA) is the preferred choice for high confidence identifications as it selects the most abundant ions by mass-to-charge ratio (m/z), thereby obtaining high-quality spectra [1][10].

Recently, tailored acquisition methods on the timsTOF instruments have resulted in high-coverage immunopeptidome profiling of biological samples. These instruments feature a dual TIMS (trapped ion mobility separation) setup, enabling parallel accumulation and serial fragmentation (PASEF)[11]. In this approach, peptides are accumulated in packages and then separated based on their ion mobility, significantly enhancing immunopeptide detection[12][13][14]. However, little attention has been given to evaluating search engines for LC-IMS-MS/MS (Liquid Chromatography Ion Mobility Mass Spectrometer) immunopeptidomics data acquired using DDA-PASEF, a hybrid methodology that integrates DDA with PASEF to achieve high-coverage data acquisition. In addition, many current programs were developed for general proteomics or tested on data from different instruments (e.g., Orbitrap).

To address this gap, we benchmarked database search algorithms for their performance in identifying immunopeptides from an optimized Thunder-DDA-PASEF acquisition strategy [14]. The search engines SEQUEST, MASCOT, and X!Tandem played a foundational role in the development of modern proteomics research. This study, however, focuses on the most current and relevant software available to date. MaxQuant [15], which uses the Andromeda search engine, is one of the most widely used tools for proteomics analysis. MSFragger from FragPipe [16] focuses on speeding up analysis by utilizing fragment ion indexing, making it ideal for searching large search spaces, such as those found in immunopeptidomics. Likewise, PEAKS [17] focuses on de novo assisted search, by generating tags for reducing the search space and yielding more robust results. The newly released MHCquant [18], built around the Comet search engine, optimizes workflows for immunopeptidomics by refining result via downstream processes such as rescoring. Analysis of the degree of consensus among the different tools serves as an indicator of the individual result’s reliability and quality. Finally, these insights will guide the selection of optimal software tools for specific research contexts.

### Experimental Procedures

#### Experimental design and Statistical Rationale

To generate the benchmarking dataset, MHC class 1 (MHC1p) and MHC class 2 peptides (MHC2p) were enriched from the JY lymphoblastoid cell line using a sequential immunoprecipitation method (See below). Four technical replicates were analyzed via three injections each (n = 12) by LC-IMS-MS employing the Thunder-DDA-PASEF method. For MHC2p analysis, the isolation polygon was modified to accommodate multiply charged ions resulting from longer peptides in the multiply charged ion cloud.

The JY MHC1p and MHC2p datasets were processed using PEAKS X Pro, PEAKS 11, FragPipe, MaxQuant, and MHCquant. Machine learning-based rescoring methods have been shown to enhance immunopeptide identification from DDA-PASEF data [14][19]. Since PEAKS 11 Studio and Online, and FragPipe include built-in rescoring, these options were activated for this analysis (deep-learning boost and MSBooster [20], respectively). Additional downstream rescoring third party tools for PEAKS X Pro and MaxQuant were not employed in order to compare software performance in their native state.

The heterogeneity in search engines, setting options, and rescoring algorithms across software packages complicates uniform comparison. To mitigate this, variables were standardized where possible (see below), with remaining parameters left at default. To address the quality of matching ambiguous spectra to peptide sequences, 1% PSM FDR filtering is employed, unless otherwise stated.

#### JY cell culture

The EBV (Epstein-Barr virus) transformed human B lymphoblastoid cell line JY (RRID:CVCL_0108)[21] was cultured in RPMI-1640 medium (Sigma-Aldrich, Cat# R8758) supplemented with 10% fetal bovine serum (PAN Biotech, Cat# P40-37100, v/v), 1 mM sodium pyruvate, 2 mM glutamine and 1% of Penicillin/Streptomycin (Sigma-Aldrich, Cat# P4333, v/v). Suspension cell cultures were harvested at 220 x g for 10min and washed three times with 1x PBS prior counting and freezing at −80 °C.

#### Immunopeptide enrichment by immunoprecipitation (IP)

In brief, cell pellets (4x10^8^ cells) were defrosted and lysed by adding cold 1% CHAPS (m/v) lysis buffer in PBS, shaking on ice, and sonicating the samples in a Bioruptor (Diagenode). Pooled samples were re-distributed into four replicates (1x10^8^ cells each) and immunoprecipitated overnight at 4 °C rotating with anti-pan-HLA1 antibody (W6/32, Leinco H263) coupled to CNBr activated beads (Cytivia). A second immunoprecipitation of the unbound fraction was performed with anti-HLA-DR antibody (L243, Leinco H261) coupled beads instead. Beads were washed 2 times with 1xPBS and once with H2O, followed by seven iterations of elution from beads with 200 µl 0.2% trifluoroacetic acid (TFA) shaking on ice. Subsequently samples were ultrafiltrated in tubes with 10 kDa molecular weight cut off (MWCO) (Vivacon 500, Hydrosart) and desalted on Oasis HLB plate (Waters) using an elution buffer of 35% acetonitrile (ACN), 0.1% TFA. Finally, samples were lyophilized and dissolved in 15 µl 0.1% formic acid (FA) shortly before LC-MS/MS analysis.

#### LC-MS/MS analysis

Desalted peptides were analyzed by nanoLC-MS using a nanoElute coupled to a timsTOF-Pro-2 mass spectrometer using Thunder-DDA-PASEF for MHC1p [14], with adaptations for MHC2p. Chromatographic separation was performed in a C18 reversed-phase column (Aurora 25 cm x 75 µm ID, 120 Å pore size, 1.7 µm particle size, IonOpticks, Australia). The samples were injected directly into the analytical column and separated using a 47-minute gradient, increasing the proportion of phase B (ACN with 0.1% FA (v/v)) to phase A (water with 0.1% FA (v/v)). The gradient started at 2% B, increased to 17% within 23 min, then to 25% in 11.5 min, to 37% in 3.8 min, and to 95% in 3.8 min, and finished with a wash step of 4.9 min at 95% B. Ionization took place using a Captive Spray source, with a capillary voltage of 1600 V, dry gas at 3.0 L/min, dry temperature at 180 °C, and TIMS-in pressure of 2.7 mBar. The MS was piloted using Compass Hystar(v5.1) and timsControl (v.4.0.5) (Bruker). The m/z acquisition range was set at 100 to 2000 Th, and the high-sensitivity detection mode was activated. TIMS was set to a range of 0.65 to 1.75 Vs/cm^2^ with accumulation and ramp time of 300 ms. The number of MS2 frames per cycle was set to three with an overlap of one. The fragmentation intensity threshold was set at 1000 and the target intensity at 20,000. For MHC1p, the Thunder isolation polygon was used to select precursors for fragmentation within the 1/K_0_ vs. m/z range of peptides with 8 to 13 amino acids, including singly and multiply charged ions [14]. For MHC2p, the isolation polygon was modified to include all multiply charged ions and singly charged ions above 550 m/z.

#### RNA-sequencing

RNA was extracted from 6 x 10^6^ JY cells using the AllPrep DNA/RNA Mini kit (QIAGEN, Cat# 80204) and treated with DNase (Invitrogen, Cat# AM 1907). RNA-seq libraries were prepared using Illumina’s TruSeq Stranded mRNA Library Prep Kit. Libraries were sequenced with the Illumina NovaSeq 6K PE 100 S4 technology.

#### Custom databases

Unless otherwise stated, the standard reference database used is the UniProt human reference proteome database (retrieved from February 2020) extended by concatenating common contaminant sequences.

For the *Arabidopsis* entrapment strategy, the final search database was generated by concatenating the standard reference database extended by the UniProt *Arabidopsis thaliana* (UP000006548) reference database (retrieved in June 2023).

For database size comparison, an updated UniProt human reference proteome was used (retrieved in June 2023). Furthermore, additional protein sequences were included based on RNA-seq data. Total RNA sequencing data from JY cells were quality controlled by Adapter trimming using TrimGalore (0.6.10) [22] and duplicate removal using clumpify (BBMap 39.01) [23]. The reads are aligned using STAR (2.7.10b)[24] with default settings on hg38 assembly and then summarized using featureCounts (Subread 2.0.6) [25] based on UCSC annotation file retrieved from June 2023. Transcripts (including isoforms) with any aligned reads are included in the database.

*De novo* transcriptome assembly was done in Galaxy. Reads were adapter-trimmed using Trimmomatic (Galaxy Version 0.38.1) [26] and duplicates were not removed. rnaSPAdes (Galaxy Version 3.15.4) [27] on paired-end reads with default settings was used for *de novo* transcriptome assembly. Coding sequences were predicted using four softwares: transdecoder (Galaxy Version 5.5.0) [28], Borf (v1.3.0) [29], GeneMarkS-T [30]and CodAN (v1.2) [31], with default settings and translated. The same set of contaminants are always included.

#### DDA database search

Raw MS data files from timsTOF (Bruker) mass spectrometer were loaded into the software as individual samples if possible (PEAKS X Pro and PEAKS 11 Online). Option to set different Bioreplicates (FragPipe) and Experiments (MaxQuant) were used and Match between runs was disabled. Unless otherwise stated, the Precursor Mass Tolerance and Fragment Mass Tolerance were set to 15ppm and 0.03 da respectively. The peptide length was set to 7-25 peptides. For PEAKS X Pro this was not possible and results were manually filtered for 7-25 length peptides. Peptide mass range and charges were set to 200-5000 and 1-4 respectively (FragPipe, MaxQuant and MHCquant). No fixed modification was set and variable modifications includes Oxidation (M), Acetylation (Protein N-term) and Cysteinylation, with a maximum of three variable modifications on peptides. Any build-in rescoring algorithm was kept on (PeptideProphet for FragPipe, Deep learning boost for PEAKS 11 Online and percolator/deeplc/ms2pip for MHCquant). Enzyme was set to none and digestion mode to unspecific/nonspecific. No extra contaminant Fasta file was provided (however, MaxQuant building contaminant files was kept in as default).

Unless otherwise stated, the Protein and Peptide FDR were set to 1 or 100%, but the PSM FDR was left at 0.01 or 1% for all but PEAKS X Pro (which doesn’t allow this setting), in which case the 1% PSM FDR was calculated manually (see statistics section). When retrieving all unfiltered candidate PSMs, PSM FDR was also set to 1 or 100%.

For FragPipe, when specifying different Bioreplicates, a file of the combined PSM result is not generated. This is generated by manually concatenating all psm.tsv files in each “exp” folder together. Unique peptide sequences are obtained by grouping peptide sequences together and keeping the entry with the highest “PeptideProphet.Probabilty” value. In the same way, a file containing all unique peptidoforms can be generated by using the “Modified Peptide” column instead of the “Peptide” column.

The identification scores used for PEAKS X Pro and PEAKS 11 Online is “X.10LgP” (higher is more confident), for FragPipe is “PeptideProphet.Probability” (higher is more confident), for MaxQuant “PEP” is used (lower is more confident) and MHCquant is “score” (lower is more confident).

#### Peptide binding prediction

Peptide binding prediction was done using netMHCpan version 4.1 [32] for MHC class 1 and netMHCIIpan version 4.1 [33] for MHC class 2 peptides. For JY cells, the MHC class 1 alleles HLA-A:02:01, HLA-B:07:02 and HLA-C:07:02 were used with a length between 8-13 amino acids. For MHC class 2 DRB1:04:04, DRB1:13:01, DRB3:01:01 and DRB4:01:03 were selected with a length of 9-22 amino acids. The rank threshold for determining strong and weak binders was kept at default.

#### Entrapment strategy

Database search analysis was conducted as described in the DDA database search section, with the exception of employing the entrapment database and omitting FDR filtering. Subsequently, results were manually filtered to achieve a 1% peptidoform FDR. Matches exclusively associated to decoy accessions were removed. Target hits were defined as peptide hits corresponding to any protein accessions from the standard reference database, while Arabidopsis false positives were classified as identified peptides associated exclusively to the Arabidopsis proteome. For comparative analysis, with the exception of MHCquant, target-decoy results were generated using an analogous approach, with decoy hits serving as false positives and excluding any matches exclusively linked to Arabidopsis protein accessions. MHCquant does not return decoy sequences and thus its 1% peptide FDR filtered results are used instead. Due to MHCquant including PTMs in its calculation its results are analogous to our definition of peptidoform.

#### Statistics

The upset plot of shared peptides between software was generated using ComplexUpset (version 1.3.3).

Amino acid bias is analyzed with the online tool Composition Profiler [34]. The background Sample is the aggregate of the identified peptides from all software without removing duplicates. The individual results from each software is compared against this background sample. Significance threshold is set at 0.005 with other settings kept at default.

Spearman correlation coefficient was calculated by ranking the peptides based on the value used to determine their identification confidence (or peptide binding rank in case of binding prediction), unless otherwise stated. Rankings are ordered in a confidence descending order. The pairwise plot is generated using GGally (version 2.2.1).

For software that did not provide built-in PSM FDR filtering capabilities, manual FDR calculation was performed. This process utilized the unfiltered PSM files generated by the software. FDR was calculated as follows:

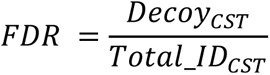

Where 𝐷𝑒𝑐𝑜𝑦_𝐶𝑆𝑇_and 𝑇𝑜𝑡𝑎𝑙_𝐼𝐷_𝐶𝑆𝑇_is the number of decoy and total identification remaining after filtering at confidence score threshold 𝐶𝑆𝑇 respectively. The 1% PSM FDR is determined by calculating the FDR at successive confidence score thresholds, moving from the most to least confident scores, and identifying the point where the FDR ≤ 0.01.

The number of unique 9mers is found by splitting all sequences into all possible 9mers and filtering out duplicates.

## Results

### A note on Peaks Studio 11 and Peaks Online 11

Bioinformatics Solutions Inc. released PEAKS Studio 11 and PEAKS Online 11 in 2023, succeeding PEAKS X (Pro). Both use identical algorithms, with PEAKS Online 11 optimized for high-throughput processing via multiple nodes [35]. We confirmed that both produce identical results given the same inputs and settings. Only minor variations were observed when activating the deep learning boost function (Supplementary Material 1), attributable to the inherent randomness of neural network models across different hardware. Due to this redundancy, only PEAKS 11 Online results were included in further analysis.

### The choice of search engine affects peptide identification from timsTOF Pro immunopeptidomics experiments

Software performance was primarily evaluated by the number of unique peptides identified at 1% PSM FDR. Here, we refer to peptides as distinct amino acid sequences and peptidoforms as PTM modified peptide variants.

Results (Figure 1A) show that four of the five software demonstrate comparable number of identified peptides. PEAKS 11 Online identified the highest number of peptides with an increase of 6-11% for both MHC class 1 and 2 peptides and peptidoforms, compared to PEAKS X Pro and FragPipe. This is followed closely by MHCquant, which saw another reduction of 6-9% compared to PEAKS X Pro and FragPipe. MaxQuant identified the fewest peptides and peptidoforms, finding 60% (MHC class 1) to 50% (MHC class 2) less compared to MHCquant.

**Figure 1.**
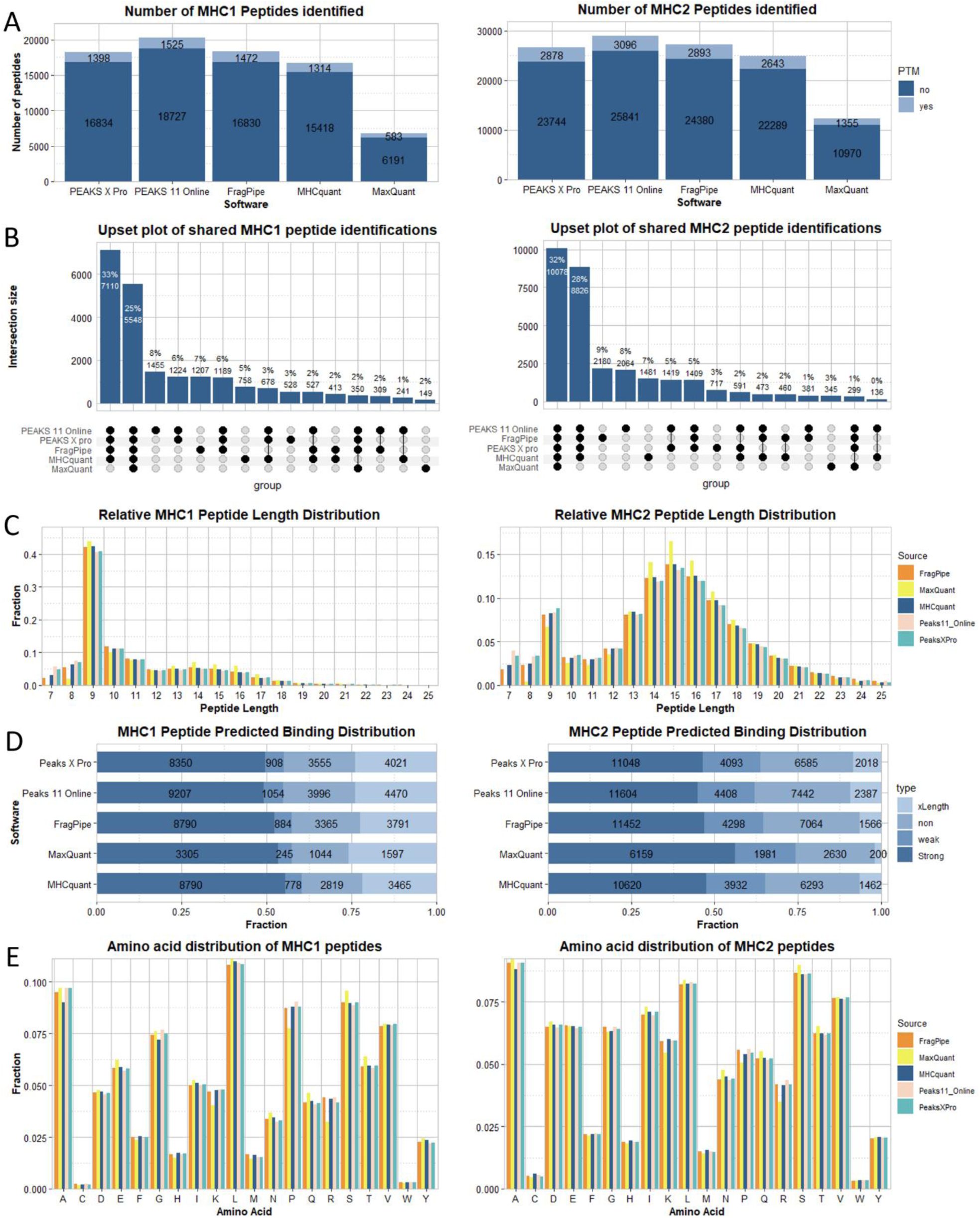
Comparative Analysis of Database Search Engine Results. MHC class 1 results are in the left column and MHC class 2 results are in the right column. A) Column chart showing the number of unique peptides identified by each software. Peptidoforms are included in lighter blue. B) Upset plot showing the intersection size of shared peptide sequences, ordered by intersection size. Only groupings resulting in an intersection size larger than 1% are included. C) Column chart showing the relative length distribution of identified peptides for each software. D) Bar chart showing the result of netMHCpan (left) and netMHCIIpan (right) binding prediction results for the result of each software, showing the fraction of strong binders (Rnk_EL ≤ 0.05% MHC class 1 and ≤ 1% class 2), weak binders (Rnk_EL > 0.05% and ≤ 2% for MHC class 1 and > 1% and ≤ 5% for class 2), non-binders (Rnk_EL > 2% for MHC class 1 and > 5% for class 2) and peptides outside the expected length (false length). E) bar chart summarizing the amino acid composition of identified peptides for each software.

To assess identification consistency across software, we evaluated shared peptide sequences (Figure 1B). 25% of MHC1p and 32% of MHC2p were identified by all software. Excluding MaxQuant, this increases to a total of 58 and 60%, respectively. MaxQuant results show the lowest absolute overlap with other software, primarily due to its lower number of identifications rather than identifying different peptides. Only between 2-10% of identified peptides are software specific, indicating general consensus in identifications.

### The peptide length distribution is largely conserved between search engines

Beyond sensitivity, assessing peptide identification biases is crucial for validating software performance. In immunopeptidomics, the length distribution of identified peptides serves as a key indicator of immunopeptide enrichment but also identification performance. Indeed, MHC1p typically have a length of 8 to 13 amino acids with a large predominance of 9-mers[3], while MHC2p extend up to 25 amino acids with a length distribution peaking at 15-mers (in humans)[36].

The length distribution is largely consistent across software packages (Figure 1C). The enrichment of 9-mers in the MHC class 1 and length distribution of MHC class 2 samples aligns with expectations and suggests successful immunopeptide enrichment and identification. The presence of 9-mers in the MHC class 2 sample may indicate a co-enrichment of MHC class 1 peptide complexes during immunoprecipitation [37]. However, MaxQuant demonstrates reduced efficiency in identifying peptides shorter than 9 amino acids. For MHC class 2, this bias is further evidenced by increased identification of longer peptides.

### MHC Binding Prediction indicates a Yield vs. Specificity Trade-off

Complementing on our analysis of peptide length distributions, we further evaluated software accuracy by leveraging predicted MHC binding affinity. This approach allows for a more comprehensive assessment of software performance in identifying biologically relevant peptides.

We used netMHCpan [32] and netMHCIIpan[33] to predict binding affinities for identified peptides (Figure 1D). The HLA alleles used are described in the Methods section. For MHC class 1 peptides, PEAKS X Pro and 11 showed similar fractions of predicted binders (0.550 and 0.548 respectively). The other three software packages demonstrated slightly higher fractions, with MHCquant performing best (0.621). For MHC class 2, MaxQuant showed the highest fraction of predicted binder (0.742). The other five software packages performed comparably. PEAKS 11 has slightly lower proportion of predicted binders, but it has the highest total number of predicted binders (10261 MHC1p and 16012 MHC2p), followed by FragPipe (9674 MHC1p and 15750 MHC2p). While a higher proportion of binders indicates better enrichment of MHCp, the detection of non-binders suggests higher sensitivity, as non-binders and co-enriched peptides with lower spectra quality are identified.

### Amino acid composition is largely conserved between search engines

Parker *et al* (2021) previously reported that search engines may prioritize different peptides based on their amino acid composition, which would introduce a bias in immunopeptide identification [38]. To evaluate if this is the case with the software tested here for DDA-PASEF data, we employed Composition Profiler [34] to quantify significant variations. This software tests whether certain amino acids or properties are enriched or depleted in a dataset compared to a background population. Here, we used the aggregate of all identified peptides as background. This approach provides a robust baseline, capturing the full spectrum of identifications and reflecting software-specific biases in amino acid composition of identified peptides, independently of other factors such as binding-motif specific enrichment. It’s important to note that this method introduces another bias, as software identifying more peptides contributes more to the background population.

Analysis of the amino acid distribution of identified peptides reveals a largely conserved pattern across software packages (Figure 1E). Notably, MaxQuant results exhibit the highest number of significant amino acid deviations for both MHC class 1 and 2 results (Supplementary Material 2). Similarly, MaxQuant most frequently shows significant differences in other amino acid properties such as charge and polarity (Supplementary Material 3). These findings suggest that MaxQuant’s peptide identification algorithm introduced the most distinct biases in the amino acid composition of reported peptides compared to other software packages. It must be noted that for MHC class 1, MHCquant has nearly the same number of significantly enriched/depleted properties as MaxQuant. Finally, visualization of motifs using unsupervised Gibbs Clustering by GibbsCluster [39] showed no obvious differences. (Supplementary Material 4). Altogether, these results indicate that, despite the slight differences, all the programs identified peptides with similar MHC binding motifs.

### Peptide identification scores correlate across software

Comparing the scores of identified peptides between software is crucial for revealing discrepancies in confidence levels and ranking of matches, potentially uncovering algorithmic biases that may be hidden when only considering overlap. However, direct comparison of absolute scores is challenging due to varying scoring methods and the impact of rescoring/boosting, which often results in bimodal score distributions. Therefore, to assess result similarity, we calculated pairwise Spearman correlation coefficients (Figure 2). Results show moderate to strong correlations between software. PEAKS X pro and FragPipe show the most similar confidence levels with a spearman correlation score of 0.8, while the lowest correlation can be observed between MaxQuant and Peaks 11 Online.

**Figure 2.**
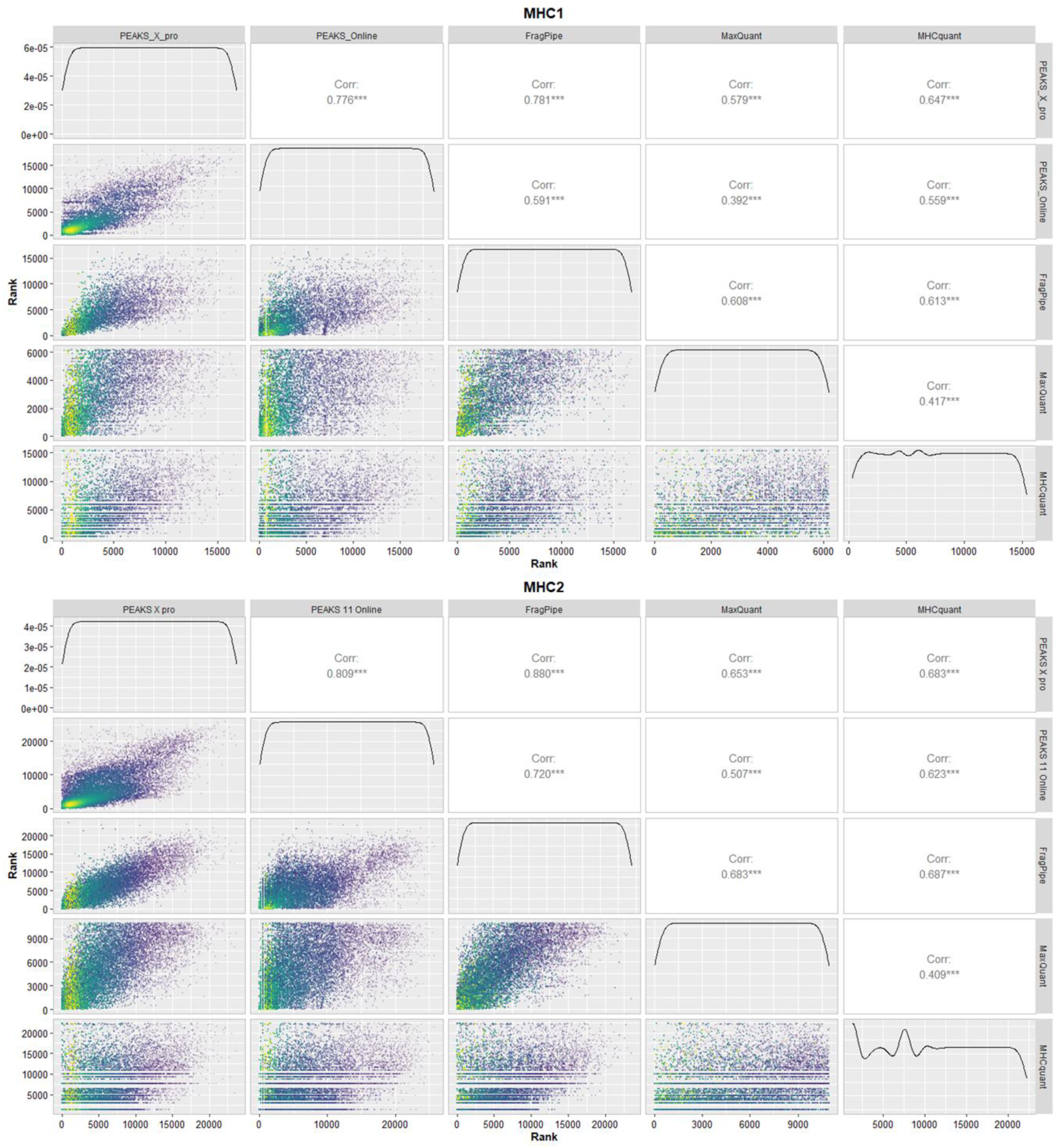
Pairwise Spearman rank correlation analysis of all software. Bottom left section displays scatterplot of ranks of shared peptides between software. Top right section returns the respective spearman rank correlation coefficient. Line plots in the center shows the rank distribution of the data. For FragPipe, hyperscore is used instead of the probability due to the lower precision caused by rounding, which leads to many shared ranks at high probabilities. Since hyperscore is used to calculate probability, they are highly correlated, making hyperscore a suitable alternative. The remaining software uses the standard values for rank calculation (i.e. “X.10LgP for PEAKS, PEP for MaxQuant and Score for MHCquant).

Next, we hypothesized that higher-affinity peptides could more likely persist through the sample preparation and accumulate for better MS identification. To evaluate if software confidence score is indicative of peptide binding strength, we evaluated the correlation between these values and NetMHCpan predicted binding rank (“Rnk_EL”). Comparison of peptides’ identification scores from each software with their netMHCpan binding rank revealed little to no correlation (Supplementary Material 5).

These findings underscore the complex interplay between peptide identification algorithms, scoring methods, and biochemical factors in immunopeptidomics, highlighting the need for cautious interpretation of software confidence scores and their relationship to immunopeptide identification.

To ensure clarity, we focus the remaining sections on MHC class 1 peptides. As of writing, MHC 1 reflects the current interest trend in the immunopeptide research community, allowing for a more manageable scope and a deeper, more consistent evaluation of search engine performance.

### *Arabidopsis* proteome entrapment sequence indicate no bias in FDR control strategy

FDR filtering using decoys helps mitigate false positives. However, unbiased implementation of this can be difficult. An effective way to assess the robustness of each software’s FDR implementation involves implementing an entrapment strategy with externally supplied entrapment sequences that act as an independent set of false positives to the software generated decoy sequences. This method helps detect potential biases from the software own FDR control implementations and provides a more comprehensive and independent evaluation of software results. Here we used *Arabidopsis thaliana* proteome sequences as entrapment sequence.

In order to compare the performance between the Arabidopsis entrapment strategy and the native target-decoy strategy, a 1% peptidoform FDR cutoff was used for both approaches. We found that a 1% PSM FDR approximates to 1.5-3.2% peptidoform FDR (and ∼5% peptide FDR if quantifiable) depending on software (Supplementary Material 6). This shows that peptidoform level FDR acts as a stricter quality control feature compared to PSM FDR.

The entrapment strategy yielded higher number of identifications at 1% peptidoform FDR for all software compared to their native target-decoy FDR control. For the software that could be tested, this gap in identification expanded as the FDR increased (Figure 3). This indicates that decoy hits are not inherently scored worse over the *Arabidopsis* false positive hits, suggesting that the native target-decoy implementations are unbiased and not overestimating FDR filtered identification.

**Figure 3.**
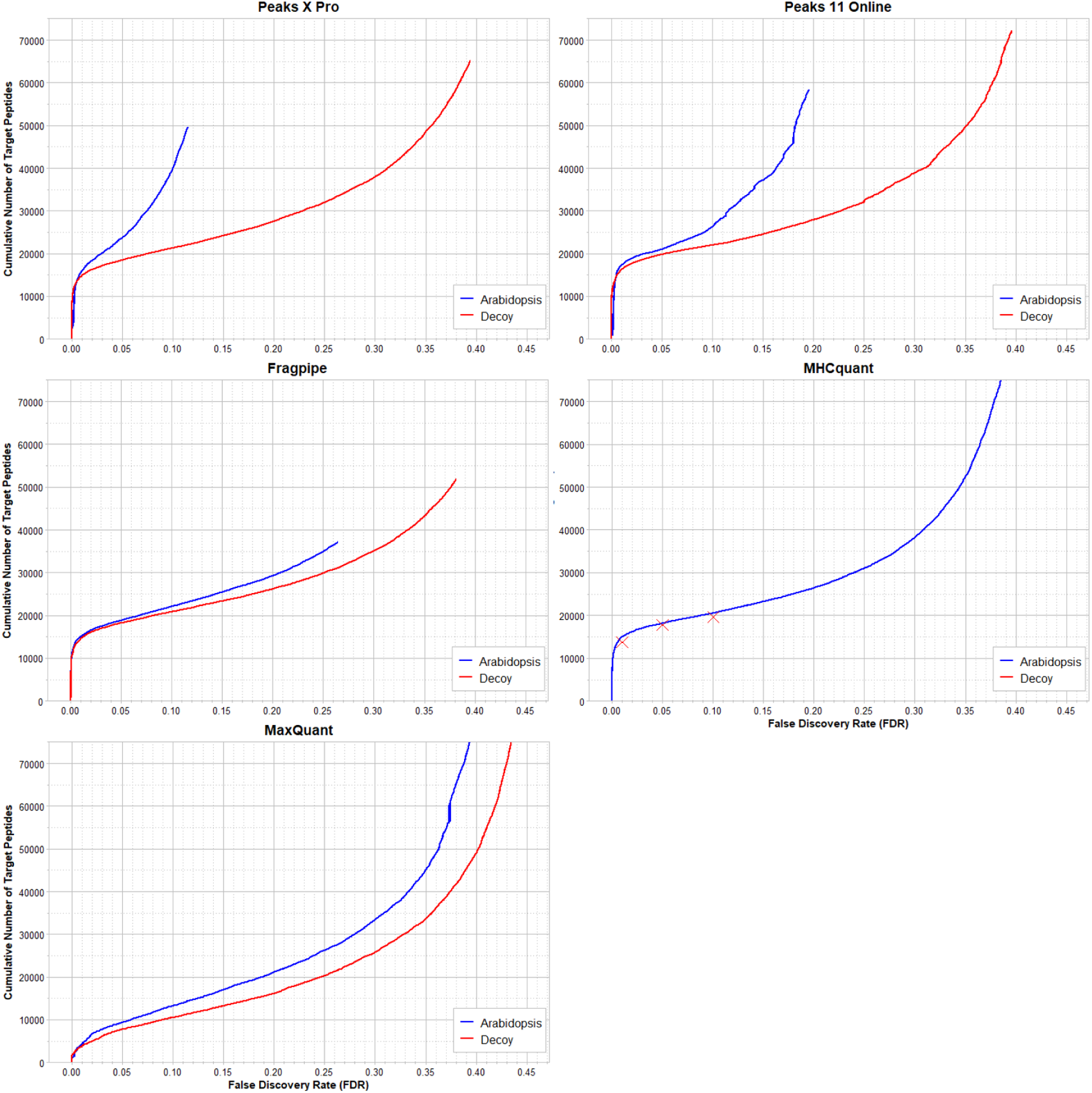
Identification performance of MHC1 peptides of different softwares using decoy and entrapment based FDR control strategy. Each graph shows the number of target peptide identified at each peptidoform FDR percentage level when using the standard decoy method (red) or the entrapment strategy (blue). Since MHCquant does not report Decoys, the software was run at 1%, 5% and 10% peptidoform FDR to obtain datapoints for comparison and displayed as “X”. y-axis limits were set to 75000.

### Database size affect peptide identification

As we have seen in the previous section, the reference database is a critical parameter in proteomics searches, affecting consistency and accuracy of peptide identifications by influencing FDR filtering and scoring. Here we further investigate this by systematically testing different database size to reveal how software adapts to changing search spaces, exposing strengths and limitations not evident with a single fixed database.

To simulate real-world research scenarios, we crafted a series of reference databases of varying sizes, based on JY cell line RNA-seq data (see Methods and Figure 4 table). This approach serves a dual purpose: it minimizes the inclusion of irrelevant sequences likely absent in our samples, while simultaneously incorporating non-canonical isoforms and accounting for edits, mutations, and variants. The latter exemplifies common use-cases in immunopeptidomics, particularly when identifying novel targets of interest.

**Figure 4.**
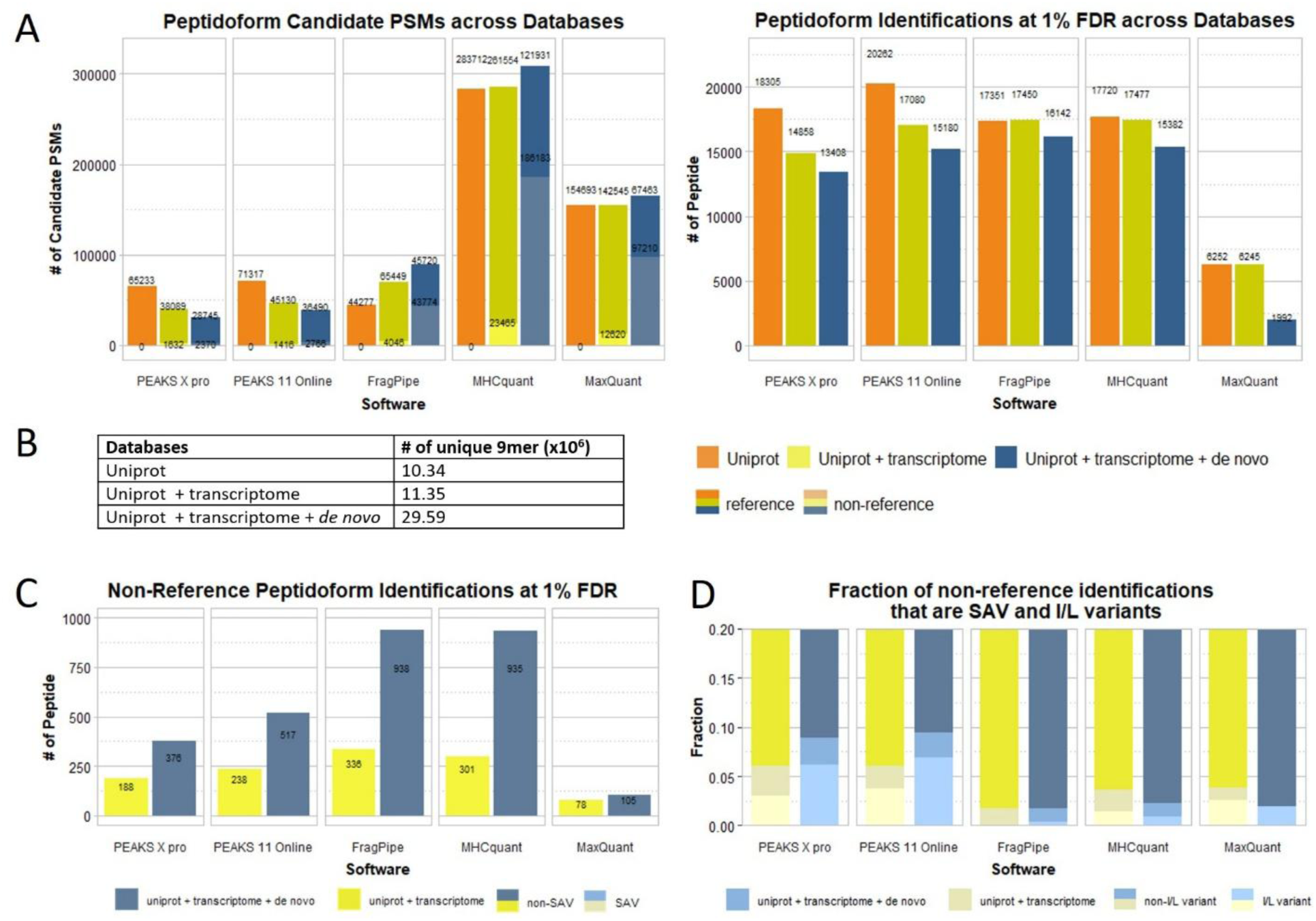
Comparison of the MHC1 identification results from different software given databases of various sizes originating from UniProt reference proteome, RNA-seq transcriptome data and de-novo transcriptome assembly. A) Grouped Stacked bar chart show the total number of identified peptidoform candidates from different software using different databases. Lighter color represents candidates not from UniProt reference database. (For the right plot, the non-UniProt reference fraction has been moved to plot C for clarity). B) Table that shows the complexity of the databases. C) Bar plot shows the number of non-reference identification at 1% FDR. D) Grouped stacked bar plot showing the fraction of non-reference peptide identifications at 1% PSM FDR filtering that are single amino acid variant (SAV) relative to the UniProt reference database and the fraction of these which are I/L variants (substitution of isoleucine with leucine or vice versa.).

As database size increases, most software packages reported increase number of candidate PSMs (at 100% PSM FDR filtering), except for PEAKS, which surprisingly exhibits a decrease (Figure 4). This unexpected trend persists despite larger databases containing a superset of sequences from smaller ones, which should theoretically allow for more comprehensive peptide-spectrum matches. The cause remains undetermined despite consultation with Bioinformatics Solutions Inc. With 1% PSM FDR filtering, the results obtained with the largest database (UniProt + transcriptome + de novo) saw the largest reduction, to the point where the number of target identifications is the lowest for all software. Furthermore, we observed that the FDR cutoff value generally becomes stricter with larger databases (Supplementary Material 7). However, this is not a perfect correlation, indicating that other aspects play a role here.

When expanding the databases, especially when including sequences derived from de novo transcriptome assembly into the database, we observed a high proportion of candidates PSMs derived from those in FragPipe, MHCquant and MaxQuant (Figure 4A lighter shaded area). However, most of these PSMs are excluded upon FDR filtering (Figure 4C).

The UniProt reference database consistently yielded the highest number of FDR filtered peptidoforms identifications for the PEAKS software, while the inclusion of the transcriptome resulted in most identifications for the other three software. The addition of sequences from the de novo transcriptome assembly data resulted in the lowest number of identifications. This indicate that addition of sequences of interest requires careful consideration to avoid overly expanding the search space, which may negatively affect the results.

DNA/RNA sequencing enables detection of peptide variants absent from reference databases. Single nucleotide variants (SNVs), the most prevalent class of genetic variants, involves a single nucleotide change that may correspond to a single amino acid substitution resulting in a single amino acid variant (SAV). We evaluated the ability of the software to distinguish SAVs through the identification of pairs of SAV and its corresponding reference. All tools successfully detected such pairs, with both PEAKS software reporting both the highest number as well as fractions of this type of variant, especially when the *de novo* transcriptome assemblies were included (Figure 4D both lighter shaded areas). However, many identified were leucine/isoleucine substitutions (Figure 4D lightest shaded area) —indistinguishable by mass—suggesting that a single spectrum may match multiple peptide candidates, depending on how results are reported. This observation raises questions about whether such cases constitute distinct identifications. In PEAKS 11 Online, 8 out of 13 and 32 out of 44 peptides are “IL variants” (for +transcriptome and +*de novo* respectively). PEAKS X Pro follows a similar trend with 5/10 and 20/29 peptides. MHCquant has 4/10 and 8/20, MaxQuant has 2/3 and 2/2 and FragPipe has 0/5 and 3/16 IL variants. From this, PEAKS software should be the method of choice when “I/L variant” detection is relevant.

## Discussion

We compared five software packages based on four algorithms for immunopeptide identification from the optimized Thunder DDA-PASEF LC-MS data acquired in a timsTOF Pro2 mass spectrometer. We demonstrate that all evaluated software packages can identify the most evident subset of immunopeptides. PEAKS 11 with the build-in rescoring algorithm exhibited superior performance in immunopeptidome identification under standard operating conditions. FragPipe emerged as a viable non-commercial alternative, yielding comparable identification results.

Software selection depends on specific research needs. Although PEAKS 11 Online identified the highest number of peptides, its performance declined with increasing database size, potentially limiting its efficacy for analyses involving extensive reference databases. Correspondence with the PEAKS software staff revealed that this performance degradation may stem from the presence of similar sequences in larger databases. They caution against using reference databases containing numerous similar sequences, such as those incorporating novel SNPs and isoforms, as this may compromise their software’s effectiveness^1^. In such cases, FragPipe is a good choice. On the other hand, PEAKS return the most I/L variant peptides, which may be important for studies emphasizing such targets.

The PEAKS 11th generation software offers an immunopeptide-specific workflow that employs a machine learning model tailored to immunopeptides. This particular approach was not incorporated in our analysis. Although this is a limitation of our study, here we focused on evaluating database search engines. Future research may benefit from a thorough examination of this workflow’s performance and potential applications in immunopeptide analysis.

Machine learning-based rescoring functions have been shown to improve peptide identification under FDR filtering [40] in MaxQuant [19] and Peaks X Pro [14], which is also seen from our results comparing Peaks X Pro and Peaks 11 Online. The entrapment strategy demonstrated unbiased target-decoy performance across all software, including those utilizing deep learning algorithms. This finding is significant, addressing concerns about potential overfitting in deep learning-based methods. However, caution is warranted when using deep learning approaches due to inherent randomness, which may affect reproducibility. Thorough evaluation of deep learning models remains essential to identify potential biases that may impact result integrity.

Our findings contribute to the optimization of immunopeptidome analysis workflows and aid in software selection for future studies in this field. Especially low quantity targets can prove challenging for identification, making the correct choice of bioinformatics analysis essential. In the future we look towards data-independent acquisition (DIA) strategies, as search engines mature and AI driven technology are incorporated. DIA allows for the simultaneous acquisition of all precursor ions within a specified mass range, improving the depth and comprehensiveness of proteomic analysis, but at the same time generates more complex spectra that require much more sophisticated processing and bioinformatic analysis.

### Database search software versions

Peaks X Pro: 10.6 build 20201221

Peaks Online 11: September 2023 version

Peaks Studio 11: build 20230202

FragPipe: 18.0/ 3.5/ 4.4.0 (FragPipe/MSFragger/Philosopher) – FragPipe 21.2-build03 was used for database size comparison.

MaxQuant: 2.4.4.0 MHCquant: 2.6.0dev

### CRediT authorship contribution statement

YC: Conceptualization, Methodology, Formal Analysis, Writing – Original Draft, Visualization, Software. AP, AA and RP: Investigation. DGZ Conceptualization, Supervision, Writing – Review & Editing. ST: Funding acquisition, Supervision, Writing – Review & Editing.

## Supporting information

Suppemental Material

## Acknowledgements

We acknowledge the support of the Immunopeptidomics Platform of HI-TRON Mainz.

We would like to acknowledge Lucas Kleinort (HI-TRON, Mainz, Germany) for his technical assistance. We also thank the Sequencing Core Facility at the German Cancer Research Center (DKFZ) for providing excellent sequencing services.

## Data availability

The mass spectrometry immunopeptidomics and proteomics data will be deposited to the ProteomeXchange Consortium [41] via the jPOSTrepo partner repository [42].

## Transcriptomic Data

RNA-seq data will be shared in Gene Expression Omnibus at NCBI [43].

## Abbreviations

DDA: Data Dependent Acquisition
DIA: Data Independent Acquisition
FDR: False Discovery Rate
MHC: Major Histocompatibility Complex
MHC1p: Major Histocompatibility Complex 1 peptide
MHC2p: Major Histocompatibility Complex 2 peptide
MS: Mass spectrometry
PASEF: Parallel Accumulation and Serial Fragmentation
PSM: Peptide-Spectrum Match
SAV: Single Amino acid Variants
SNV: Single Nucleotide Variants
TIMS: Trapped Ion Mobility Separation

## Footnotes

^1^Kristina Jurcic (Bioinformatics Solutions Inc.), oral communication, 26^th^ April 2024.

