## Supplementary material for "Benchmarking Software for DDA-PASEF Immunopeptidomics": Suppemental Material

#### Supplementary Material 1

| Settings | MHC1 | MHC2 | MHC1 Boost | MHC2 Boost |
|---|---|---|---|---|
| 6-45 length, 2 max mods, no boost | 0 19154 0<br>Studio Online | 0 27306 0<br>Studio Online | 138 21545 210<br>Studio Online | 300 29956 261<br>Studio Online |
| 20ppm precursor, 0.05da, no boost | 0 17166 0<br>Studio Online | 0 25504 0<br>Studio Online | 172 18903 141<br>Studio Online | 237 27716 233<br>Studio Online |
| 6-45 length, no boost | 0 19165 0<br>Studio Online | 0 27556 0<br>Studio Online | 221 21606 212<br>Studio Online | 308 30145 359<br>Studio Online |

*Supplementary Material 1 Venn Diagram to compare the number of identification and overlap between PEAKS 11 Online and PEAKS 11 Studio using various settings and with or without deep learning boost*

#### Supplementary Material 2

| MHC1 | Peaks 11 Online | Peaks X pro | FragPipe | MaxQuant | MHCquant |
|---|---|---|---|---|---|
| Ala | Not significant. | Not significant. | Not significant. | Not significant. | Depleted. |
| Arg | Enriched. | Not significant. | Enriched. | Depleted. | Not significant. |
| Asn | Depleted. | Not significant. | Not significant. | Enriched. | Not significant. |
| Asp | Not significant. | Not significant. | Not significant. | Not significant. | Not significant. |
| Cys | Not significant. | Not significant. | Not significant. | Not significant. | Not significant. |
| Gln | Not significant. | Not significant. | Not significant. | Enriched. | Not significant. |
| Glu | Not significant. | Not significant. | Not significant. | Enriched. | Not significant. |
| Gly | Enriched. | Not significant. | Not significant. | Not significant. | Depleted. |
| His | Not significant. | Not significant. | Not significant. | Depleted. | Not significant. |

| | | | | | |
|---|---|---|---|---|---|
| Ile | Not significant. | Not significant. | Not significant. | Not significant. | Not significant. |
| Leu | Not significant. | Not significant. | Not significant. | Not significant. | Not significant. |
| Lys | Not significant. | Not significant. | Not significant. | Depleted. | Not significant. |
| Met | Not significant. | Not significant. | Not significant. | Not significant. | Not significant. |
| Phe | Not significant. | Not significant. | Not significant. | Not significant. | Not significant. |
| Pro | Enriched. | Not significant. | Not significant. | Depleted. | Not significant. |
| Ser | Not significant. | Not significant. | Not significant. | Enriched. | Not significant. |
| Thr | Not significant. | Not significant. | Not significant. | Enriched. | Not significant. |
| Trp | Not significant. | Not significant. | Not significant. | Not significant. | Not significant. |
| Tyr | Not significant. | Not significant. | Not significant. | Not significant. | Not significant. |
| Val | Not significant. | Not significant. | Not significant. | Not significant. | Not significant. |

| MHC2 | Peaks 11 Online | Peaks X pro | FragPipe | MaxQuant | MHCquant |
|---|---|---|---|---|---|
| Ala | Not significant. | Not significant. | Not significant. | Not significant. | Depleted. |
| Arg | Enriched. | Not significant. | Not significant. | Depleted. | Not significant. |
| Asn | Not significant. | Not significant. | Not significant. | Enriched. | Not significant. |
| Asp | Not significant. | Not significant. | Not significant. | Not significant. | Not significant. |
| Cys | Not significant. | Depleted. | Not significant. | Depleted. | Enriched. |
| Gln | Not significant. | Not significant. | Not significant. | Enriched. | Not significant. |
| Glu | Not significant. | Not significant. | Not significant. | Not significant. | Not significant. |
| Gly | Not significant. | Not significant. | Not significant. | Not significant. | Not significant. |
| His | Not significant. | Not significant. | Not significant. | Not significant. | Not significant. |
| Ile | Not significant. | Not significant. | Not significant. | Enriched. | Not significant. |
| Leu | Not significant. | Not significant. | Not significant. | Not significant. | Not significant. |
| Lys | Not significant. | Not significant. | Not significant. | Depleted. | Not significant. |
| Met | Not significant. | Not significant. | Not significant. | Depleted. | Not significant. |
| Phe | Not significant. | Not significant. | Not significant. | Not significant. | Not significant. |
| Pro | Enriched. | Not significant. | Not significant. | Depleted. | Not significant. |
| Ser | Not significant. | Not significant. | Not significant. | Enriched. | Not significant. |
| Thr | Not significant. | Not significant. | Not significant. | Enriched. | Not significant. |

| | | | | | |
|---|---|---|---|---|---|
| Trp | Not significant. | Not significant. | Not significant. | Not significant. | Not significant. |
| Tyr | Not significant. | Not significant. | Not significant. | Not significant. | Not significant. |
| Val | Not significant. | Not significant. | Not significant. | Not significant. | Not significant. |

*Supplementary Material 2. Results from Composition Profiler to determine the amino acid bias of identified peptides from each software. The sum of all identifications from all software were used as background. p < 0.005 is used as significance threshold.*

#### Supplementary Material 3

| MHC1 | Peaks 11 Online | Peaks X pro | FragPipe | MaxQuant | MHCquant |
|---|---|---|---|---|---|
| Aromatic content | Not significant. | Not significant. | Not significant. | Not significant. | Not significant. |
| Charged residues | Not significant. | Not significant. | Not significant. | Depleted. | Not significant. |
| Positively charged | Enriched. | Not significant. | Not significant. | Depleted. | Not significant. |
| Negatively charged | Depleted. | Not significant. | Not significant. | Enriched. | Not significant. |
| Polar (Zimmerman) | Not significant. | Not significant. | Not significant. | Depleted. | Enriched. |
| Hydrophobic (Eisenberg) | Enriched. | Not significant. | Not significant. | Not significant. | Depleted. |
| Hydrophobic (K-D) | Not significant. | Not significant. | Not significant. | Not significant. | Not significant. |
| Hydrophobic (F-P) | Not significant. | Not significant. | Not significant. | Not significant. | Not significant. |
| Exposed (Janin) | Not significant. | Not significant. | Not significant. | Not significant. | Enriched. |
| Flexible (Vihinen) | Not significant. | Not significant. | Not significant. | Depleted. | Not significant. |
| High interface prop. (J-T) | Not significant. | Not significant. | Not significant. | Depleted. | Enriched. |
| High solvation poten. (J-T) | Not significant. | Not significant. | Not significant. | Not significant. | Depleted. |
| Frequent in alpha hel. (N) | Not significant. | Not significant. | Not significant. | Not significant. | Not significant. |
| Frequent in beta struc. (N) | Depleted. | Not significant. | Not significant. | Enriched. | Enriched. |

| | | | | | |
|---|---|---|---|---|---|
| Frequent in coils (N) | Not significant. | Not significant. | Not significant. | Not significant. | Not significant. |
| High linker propensity (G-H) | Not significant. | Not significant. | Not significant. | Depleted. | Enriched. |
| Disorder promoting (Dunker) | Enriched. | Not significant. | Not significant. | Depleted. | Depleted. |
| Order promoting (Dunker) | Not significant. | Not significant. | Not significant. | Enriched. | Enriched. |
| Bulky (Zimmerman) | Not significant. | Not significant. | Not significant. | Not significant. | Not significant. |
| Large (Dawson) | Not significant. | Not significant. | Not significant. | Depleted. | Enriched. |

#### MHC2

| | | | | | |
|---|---|---|---|---|---|
| Aromatic content | Not significant. | Not significant. | Not significant. | Not significant. | Not significant. |
| Charged residues | Not significant. | Not significant. | Not significant. | Depleted. | Not significant. |
| Positively charged | Enriched. | Not significant. | Not significant. | Depleted. | Not significant. |
| Negatively charged | Not significant. | Not significant. | Not significant. | Not significant. | Not significant. |
| Polar (Zimmerman) | Not significant. | Not significant. | Not significant. | Depleted. | Enriched. |
| Hydrophobic (Eisenberg) | Not significant. | Not significant. | Not significant. | Not significant. | Not significant. |
| Hydrophobic (K-D) | Not significant. | Not significant. | Not significant. | Enriched. | Not significant. |
| Hydrophobic (F-P) | Not significant. | Not significant. | Not significant. | Not significant. | Not significant. |
| Exposed (Janin) | Not significant. | Not significant. | Not significant. | Not significant. | Not significant. |
| Flexible (Vihinen) | Not significant. | Not significant. | Not significant. | Depleted. | Not significant. |
| High interface prop. (J-T) | Not significant. | Not significant. | Not significant. | Not significant. | Enriched. |
| High solvation poten. (J-T) | Not significant. | Not significant. | Not significant. | Not significant. | Not significant. |
| Frequent in alpha hel. (N) | Not significant. | Not significant. | Not significant. | Not significant. | Not significant. |

| | | | | | |
|---|---|---|---|---|---|
| Frequent in beta struc. (N) | Not significant. | Not significant. | Not significant. | Enriched. | Not significant. |
| Frequent in coils (N) | Not significant. | Not significant. | Not significant. | Not significant. | Not significant. |
| High linker propensity (G-H) | Not significant. | Not significant. | Not significant. | Depleted. | Not significant. |
| Disorder promoting (Dunker) | Enriched. | Not significant. | Enriched. | Depleted. | Depleted. |
| Order promoting (Dunker) | Not significant. | Not significant. | Not significant. | Enriched. | Not significant. |
| Bulky (Zimmerman) | Not significant. | Not significant. | Not significant. | Enriched. | Not significant. |
| Large (Dawson) | Not significant. | Not significant. | Not significant. | Depleted. | Enriched. |

*Supplementary Material 3 Results from Composition Profiler to determine property bias of identified peptides from each software.*

*The sum of all identifications from all software were used as background. p < 0.005 is used as significance threshold.*

**Supplementary Material 4**

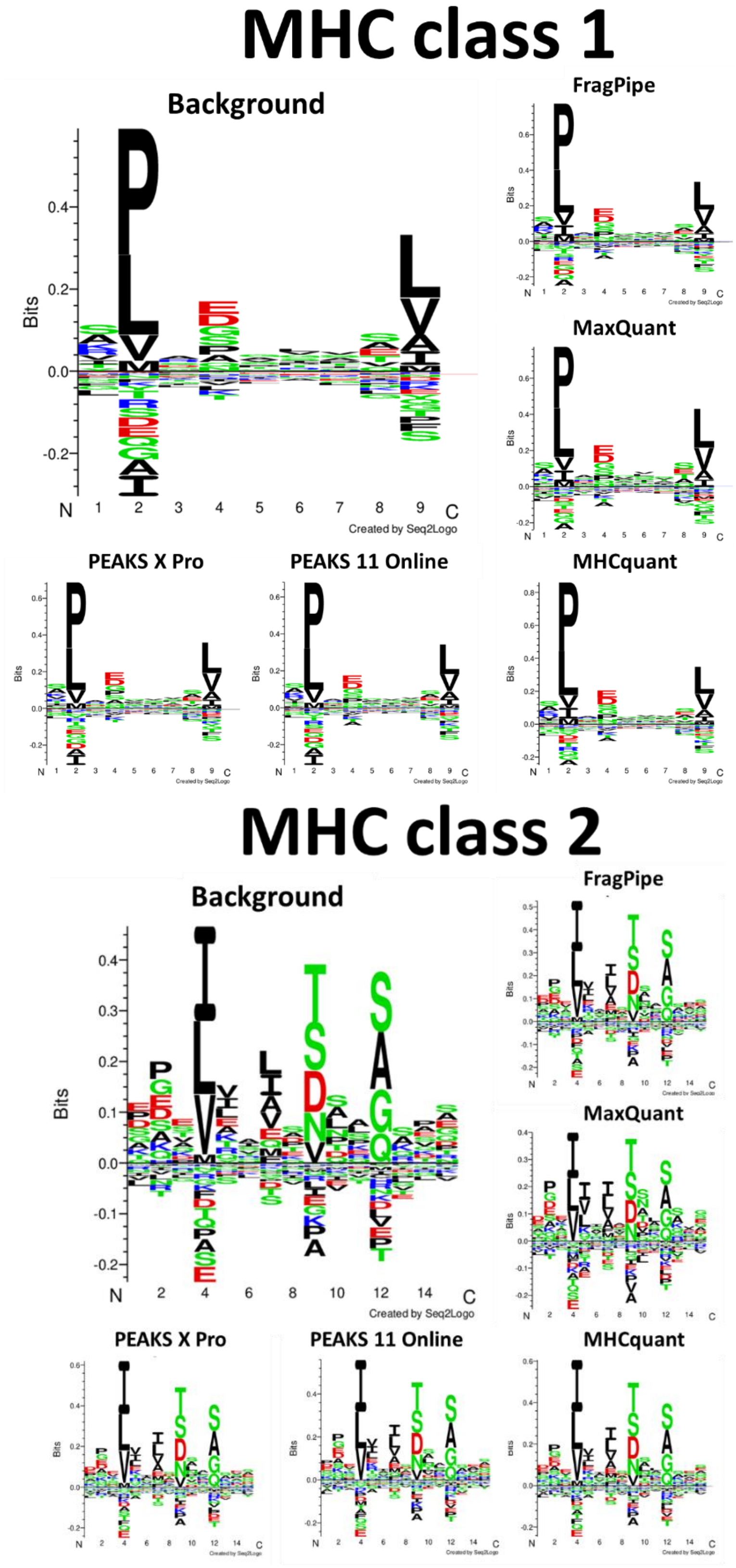

*Supplementary Material 4 Unsupervised Gibbs clustering of motifs for MHC class 1 (Top) and MHC class 2 (Bottom) peptide identifications from each software. Background is the combination of peptide identifications from all software. For MHC1, a motif length of 9 amino acids was used. For MHC class 2, 15 amino acid motif length was set.*

### Supplementary Material 5

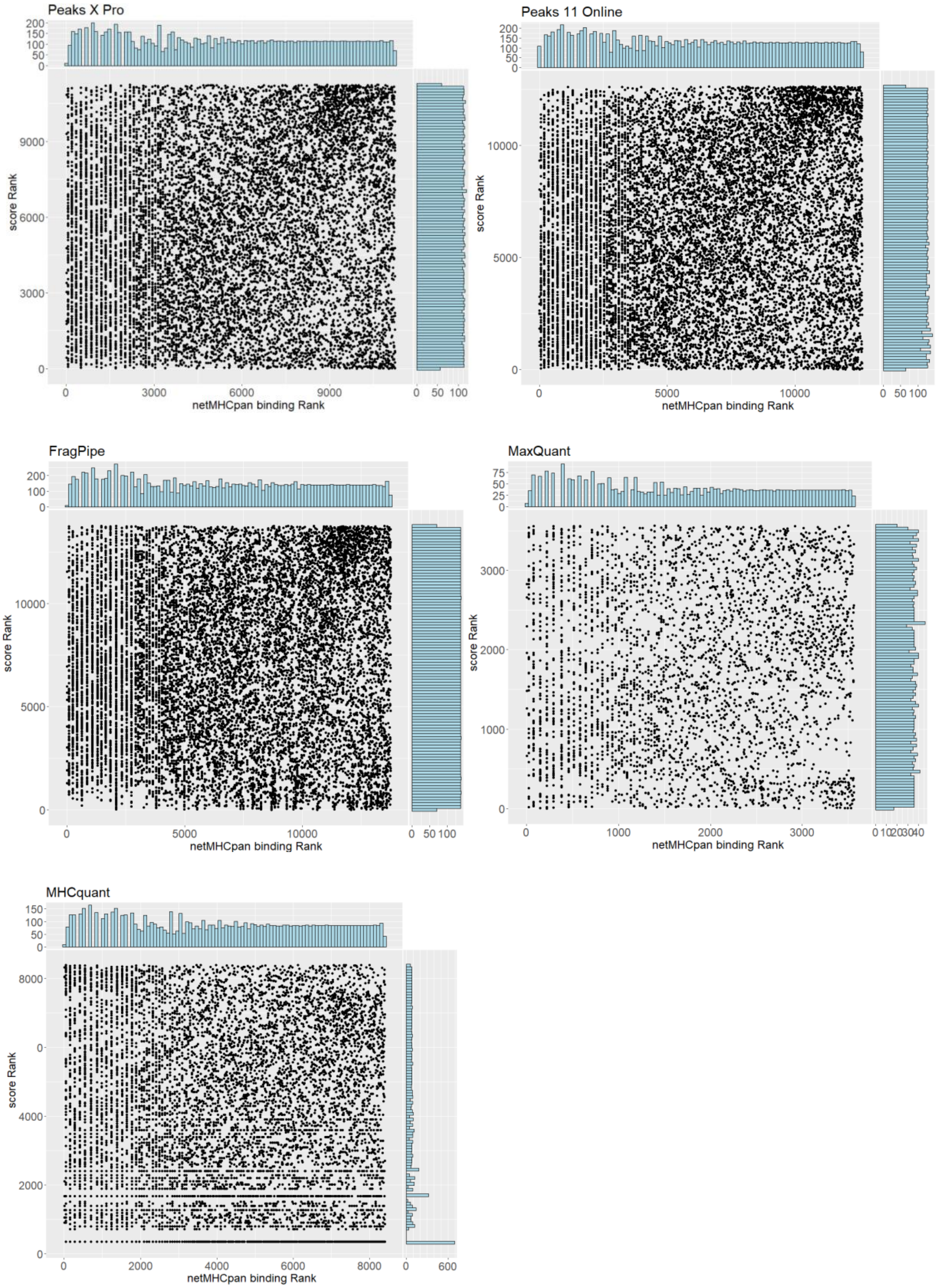

| | PEAKS X Pro | PEAKS 11 Online | FragPipe | MaxQuant | MHCquant |
|---|---|---|---|---|---|
| **Spearman correlation coefficient** | 0.120 | 0.160 | 0.0598 | 0.089 | 0.0002 |

*Supplementary Material 5 Top) Scatterplot visualizing the peptide identification confidence as a function of their score rank for each software with their predicated binding strength from netMHCpan. Bottom) Table showing the spearman correlation for each of the scatterplots*

#### Supplementary Material 6

Number of unique peptides found at different PSM and peptide and peptidoform FDR (Decoy) on standard

| MHC1 | Peaks X pro | Peaks 11 Online | FragPipe | MHCquant | MaxQuant |
|---|---|---|---|---|---|
| 1% PSM | 16834 (2.7% peptidoform) | 18727 (3.2% peptidoform FDR) | 16830 (1.5% peptidoform) | 15418 | 6191 (2.6% peptidoform) |
| 1% peptide | 13482 | 14719 | 13056 | n/a | 3812 |
| 5% peptide | 16797 | 18110 | 16531 | n/a | 7356 |
| 10% peptide | 19499 | 20091 | 18930 | n/a | 10129 |
| 1% peptidoform | 14890 | 16269 | 14482 | 13765 | 4231 |
| 5% peptidoform | 18506 | 19799 | 18226 | 17745 | 8389 |
| 10% peptidoform | 21539 | 21986 | 20832 | 19582 | 11459 |

Number of unique **peptide** above 1% **peptide** FDR when using Decoy or ARATH as false positive (value in brackets denote score cutoff)

| MHC1 | Peaks X pro | Peaks 11 Online | FragPipe | MHCquant | MaxQuant |
|---|---|---|---|---|---|

| | | | | | |
|---|---|---|---|---|---|
| Decoy on std | 13482 (21.85) | 14719 (70.71) | 13056 (0.9916) | n/a | 3812 (0.01277) |
| Decoy on mixed | 12987 (19.48) | 14345 (68.1) | 12912 (0.9840) | n/a | 2756 (0.01272) |
| Arath on mixed | 14555 (15.34) | 15802 (49.16) | 13632 (0.9750) | 13647 (0.00506) | 3792 (0.02090) |

Number of unique peptides found at different **peptide** FDR (ARATH)

| MHC1 | Peaks X pro | Peaks 11 Online | FragPipe | MHCquant | MaxQuant |
|---|---|---|---|---|---|
| 1% peptide | 14555 | 15802 | 13632 | 13647 | 3792 |
| 5% peptide | 21686 | 19333 | 17118 | 16539 | 8273 |
| 10% peptide | 36216 | 23867 | 20066 | 18748 | 11732 |

Number of unique **peptidoform** above 1% **peptidoform** FDR when using Decoy or ARATH as false positive (value in brackets denote score cutoff)

| MHC1 | Peaks X pro | Peaks 11 Online | FragPipe | MHCquant | MaxQuant |
|---|---|---|---|---|---|
| Decoy on std | 14890 (21.44) | 16269 (68.30) | 14482 (0.9907) | 13765 (0.00995) | 4231 (0.01407) |
| Arath on mixed | 15914 (16.65) | 17289 (47.68) | 15077 (0.9730) | 14944 (0.005373) | 4302 (0.02351) |

Number of unique peptides found at different **peptidoform** FDR (ARATH)

| MHC1 | Peaks X pro | Peaks 11 Online | FragPipe | MHCquant | MaxQuant |
|---|---|---|---|---|---|
| 1% peptidoform | 15914 | 17289 | 15077 | 14944 | 4302 |
| 5% peptidoform | 23671 | 21024 | 18791 | 18151 | 9327 |
| 10% peptidoform | 39894 | 26221 | 22067 | 20492 | 13189 |

*Supplementary Material 6 Tables summarizing the number of identified MHC1p (and cutoff score) from each software at various FDR and with various FDR strategies: PSM, peptide and peptidoform. Peptides are defined as sequences with the PTMs removed, while peptidoforms retain the PTMs. The database used for these results are either the standard database (std) or the Arabidopsis entrapment database (mixed). False positives used for calculating the FDR is either decoy or Arabiopsis thaliana (Arath).*

#### Supplementary Material 7

| | UniProt | UniProt + transcriptome | UniProt + transcriptome + de novo |
|---|---|---|---|
| Peaks X Pro (higher = stricter) | 15.97 | 18.06 | 18.10 |
| Peaks 11 Online (higher = stricter) | 24.38 | 24.25 | 31.21 |
| FragPipe (higher = stricter) | 0.9214 | 0.9226 | 0.9256 |
| MaxQuant (lower = stricter) | 0.035702 | 0.036796 | 0.026930 |
| MHCquant (lower = stricter) | 0.00999848 | 0.00999521 | 0.00999834 |

*Supplementary Material 7 Tables showing the 1% PSM FDR cutoff value for all software. Higher = stricter: A higher value indicates greater confidence in the identification, therefore having a cutoff at a higher value is considered stricter. Likewise for the lower = stricter.*
